# Using machine learning-based lesion behavior mapping to identify anatomical networks of cognitive dysfunction: spatial neglect and attention

**DOI:** 10.1101/556753

**Authors:** Daniel Wiesen, Christoph Sperber, Grigori Yourganov, Christopher Rorden, Hans-Otto Karnath

## Abstract

Previous lesion behavior studies primarily used univariate lesion behavior mapping techniques to map the anatomical basis of spatial neglect after right brain damage. These studies led to inconsistent results and lively controversies. Given these inconsistencies, the idea of a widespread network that might underlie spatial orientation and neglect has been pushed forward. In such case, univariate lesion behavior mapping methods might have been inherently limited in uncover the presumed network in a single study due to limited statistical power. By using multivariate lesion-mapping based on support vector regression, we aimed to validate the network hypothesis directly in a large sample of 203 newly recruited right brain damaged patients. In a single analysis, this method identified a network of parietal, temporal, frontal, and subcortical regions, which also included white matter tracts connecting these regions. The results were compared to univariate analyses of the same patient sample using different combinations of lesion volume correction and statistical thresholding. The comparison revealed clear benefits of multivariate lesion behavior mapping in identifying brain networks.

## 1. Introduction

Spatial attention and orientation is a cognitive function dominantly represented in the human right hemisphere (Corbetta et al., 2008, 2005). In correspondence, spatial neglect is one of the most common syndromes after brain injury of predominantly this hemisphere (Stone et al., 1993; Becker and Karnath, 2007; Ten Brink et al., 2017). Patients spontaneously and sustainably deviate towards the ipsilesional side, neglecting contralesionally located information or stimuli (Heilman et al., 1983; Karnath and Rorden, 2012). The anatomical basis of this core deficit of spatial neglect has been extensively investigated during the last decade by using mass-univariate lesion-behavior mapping methods (VLBM), such as VLSM (Bates et al., 2003) or NPM (Rorden et al., 2007). Heterogeneous findings were observed, causing lively controversies (for review Karnath and Rorden, 2012). In the right hemisphere, spatial neglect has been reported to be associated with parietal lesions to the inferior parietal lobule and temporo-parietal junction (Chechlacz et al., 2010; Karnath et al., 2011; Rousseaux et al., 2015), the superior and middle temporal cortex as well as the insula (Karnath et al., 2004, 2011; Committeri et al., 2007; Sarri et al., 2009; Chechlacz et al., 2010; Saj et al., 2012; Rousseaux et al., 2015) and the ventrolateral prefrontal cortex (Committeri et al., 2007; Thiebaut De Schotten et al., 2014). These cortical areas were also found to be involved in the human left hemisphere when patients show spatial neglect after a left hemisphere stroke (Suchan and Karnath, 2011). Furthermore, disrupted structural connectivity has been related to spatial neglect, including damage of the superior longitudinal fasciculus and arcuate fasciculus, the inferior occipito-frontal fasciculus, extreme capsule and the superior occipito-frontal fasciculus, as well as the middle longitudinal fasciculus (Thiebaut De Schotten et al., 2005; He et al., 2007; Urbanski et al., 2008, 2011; Karnath et al., 2009; Shinoura et al., 2009; Ciaraffa et al., 2013; Thiebaut De Schotten et al., 2014; Umarova et al., 2014; Vaessen et al., 2016; Carter et al., 2017; de Haan and Karnath, 2017).

Building on the seminal work by Watson and colleagues (1974) and Mesulam (1981), it has been concluded in review articles (Catani, 2006; Bartolomeo et al., 2007; Karnath, 2009; Karnath and Rorden, 2012; Lunven and Bartolomeo, 2017) and meta-analyses (Chechlacz et al., 2012; Molenberghs et al., 2012) that an anatomical network, also termed as ‘perisylvian network’ (Karnath, 2009; Karnath and Rorden, 2012), might represent the basis of spatial neglect. However, so far none of the previous mass-univariate lesion-behavior studies has empirically confirmed this network in a single analysis. In fact, traditional mass-univariate lesion-behavior mapping methods might be ill-suited to provide an empirical confirmation of the network hypothesis in spatial neglect. Due to the so-called ‘partial injury problem’ (Rorden et al., 2009; Sperber et al., in press), statistical power of VLBM in anatomical networks might be reduced, and false negative findings might conceal the full network. This issue has been confirmed by several simulation studies (Mah et al., 2014; Zhang et al., 2014; Pustina et al., 2018). Furthermore, the huge number of independent tests in VLBM as well as in some of the multivariate lesion behavior mapping (MLBM) implementations requires statistical control for multiple comparisons, which can further reduce statistical power. This might have contributed to the heterogeneous pattern of previous VLBM results in spatial neglect, with different studies identifying some nodes while missing others. Additional facts which can explain heterogeneous anatomical findings might be based on the specific sample characteristics in previous investigations (e.g., see Gajardo-Vidal et al., 2018). The authors showed that specific sub-sets of patients might drive significant results. However, this might be especially true in smaller studies with lower power (Lorca-Puls et al., 2018). Another influential factor are the clinical tests administered (for review, see Sperber and Karnath, 2018).

The label “spatial neglect” has often been used as an umbrella term interchangeably for a collection of various symptoms. In line with recent efforts to center on different subcomponents observed in spatial neglect, rather than aggregating them in one analysis (Vuilleumier, 2013), we here focus on only the egocentric core component of spatial neglect (see Karnath and Rorden, 2012). This core component is represented by a spontaneous and sustained deviation of eyes and head towards the ipsilesional side (Fruhmann-Berger and Karnath, 2005; Fruhmann-Berger et al., 2006; Becker and Karnath, 2010), combined with neglect of contralesionally located information or stimuli. The spatial bias can be reliably measured amongst others by traditional cancellation tasks (Rorden and Karnath, 2010) as well as a modified line bisection task (McIntosh et al., 2017). Further spatial and non-spatial symptoms that have been described in neglect patients (e.g. Binder et al., 1992; Husain et al., 1997; Barton and Black, 1998; Ferber and Karnath, 2001; Azouvi, 2002; Husain and Rorden, 2003; Verdon et al., 2010; Sperber et al., 2016; McIntosh et al., 2017) were not intended to be covered by the present study.

Multivariate lesion behavior mapping (MLBM) appears to be particularly suitable to identify neural correlates of behavior organized in networks (Smith et al., 2013; Mah et al., 2014; Zhang et al., 2014; Yourganov et al., 2015; Zavaglia and Hilgetag, 2016; Pustina et al., 2018; for review Karnath et al., 2018). Two recent studies investigated the neural correlates of spatial neglect with MLBM, one using support vector machines in a sample of 140 right hemisphere stroke patients (Smith et al., 2013), the other a game theoretical approach in a small sample of only 25 (and even less for subtests) right hemisphere patients (Toba et al., 2017). Both studies had limitations. The two approaches were constrained to the investigation of only a few brain regions in a single analysis at once, in other words, these approaches did not provide a voxel-by-voxel analysis of the lesion pattern throughout the brain. Moreover, regions in such region-based approaches can differ both from the relevant functional parcellation of the brain and the typical anatomy of stroke lesions and thus might have failed to capture relevant brain regions. In contrast, Support Vector Regression based Multivariate Lesion-Symptom Mapping (SVR-LSM) utilizes the full voxel-wise whole brain information independently of an a priori region of interest selection (Zhang et al., 2014). Different groups have recently validated and tested this approach (Zhang et al., 2014, DeMarco and Turkeltaub, 2018, Sperber et al., in press). We use this approach here to explore spatial neglect in a large sample of 203 newly recruited, right hemisphere damaged patients. Our main objective was to use this method to delineate the anatomical network assumed to underlie spatial neglect and to compare its outcome to traditional mass-univariate analyses by performing VLBM analyses on the same patient sample.

## 2. Materials and Methods

### 2.1. Subjects

Neurological patients consecutively admitted to the Center of Neurology at Tuebingen University were screened for a first ever right-hemisphere stroke. Patients with a left-sided stroke, patients with diffuse or bilateral brain lesions, patients with tumors, as well as patients in whom MRI or CT scans revealed no obvious lesions were not included. In total 203 patients were recruited. None of these patients were included in any of our previous studies addressing the anatomy of spatial neglect (Karnath et al., 2001, 2004, 2011; Smith et al., 2013). Therefore, they represent an independent, new sample. Table 1 gives the demographic and clinical data. All subjects provided written informed consent and the study was conducted in accordance with the ethical guidelines from the revised Declaration of Helsinki and in accordance with relevant guidelines and regulations.

**Table 1:**
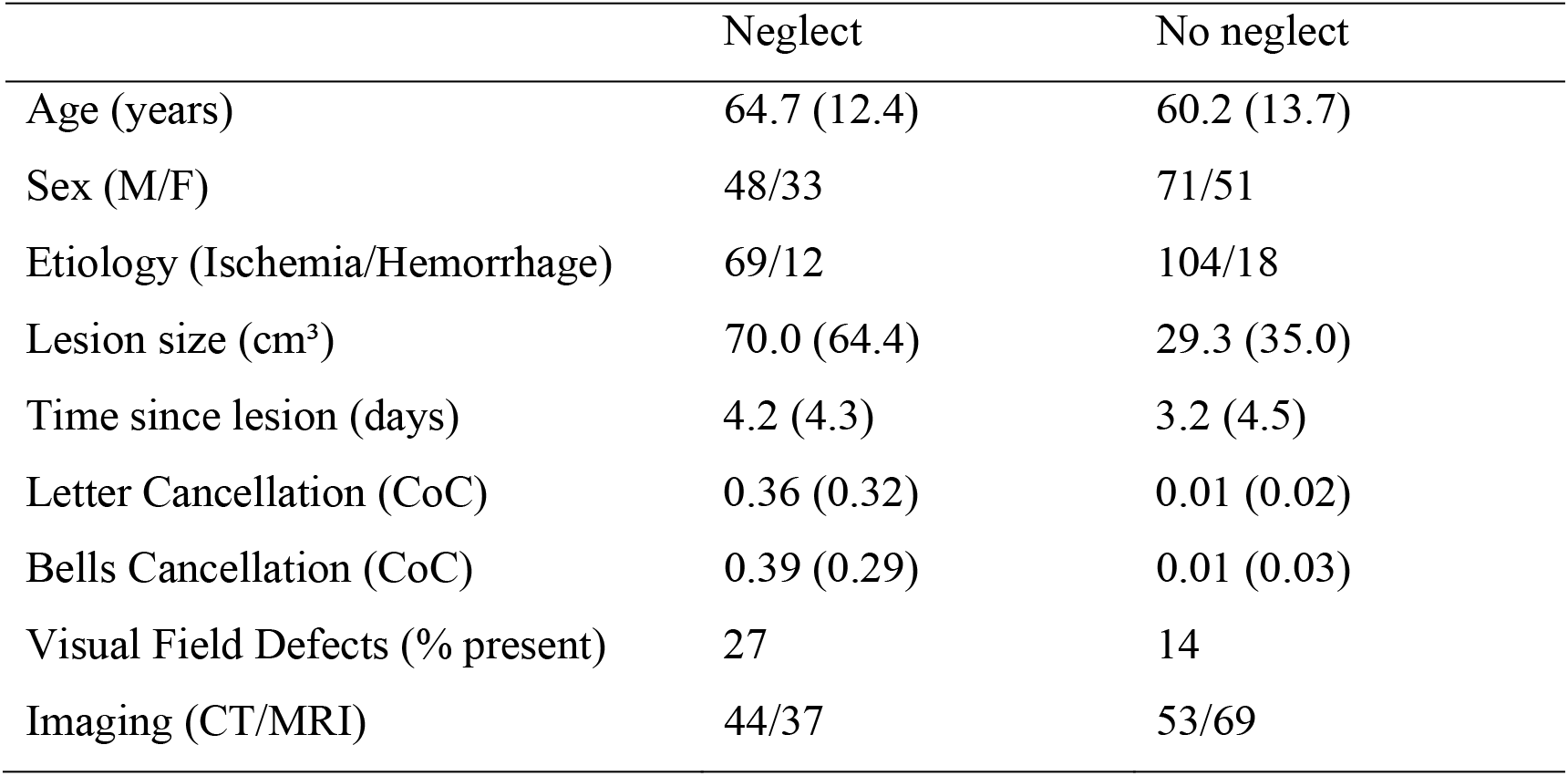
Demographic and clinical data of the 203 patients included. For this table we determined whether a CoC (= Center of Cancellation) score was in the pathological range; cut-offs were set at >.081 for the Bells Cancellation Task and >.083 for the Letter Cancellation test (cf. Rorden and Karnath, 2010). In order to assign the diagnosis of spatial neglect, patients had to present a pathological test score in at least one of the two cancellation tests. Using this criterion, 81 (40%) were classified as exhibiting spatial neglect while 122 (60%) did not exhibit neglect. Data are represented as mean (SD).

|  | Neglect | No neglect |
| --- | --- | --- |
| Age (years) | 64.7 (12.4) | 60.2 (13.7) |
| Sex (M/F) | 48/33 | 71/51 |
| Etiology (Ischemia/Hemorrhage) | 69/12 | 104/18 |
| Lesion size (cm<sup>3</sup>) | 70.0 (64.4) | 29.3 (35.0) |
| Time since lesion (days) | 4.2 (4.3) | 3.2 (4.5) |
| Letter Cancellation (CoC) | 0.36 (0.32) | 0.01 (0.02) |
| Bells Cancellation (CoC) | 0.39 (0.29) | 0.01 (0.03) |
| Visual Field Defects (% present) | 27 | 14 |
| Imaging (CT/MRI) | 44/37 | 53/69 |

### 2.2. Behavioral examination

The interval between stroke-onset and neuropsychological examination was maximally 25 days (mean = 4.37 days, SD = 4.04). The following neuropsychological tests were performed: Letter Cancellation Task (Weintraub and Mesulam, 1985) and Bells Test (Gauthier, Louise Dehaut, Francois Joanette, 1989). These two tests were presented on a horizontally oriented 21 x 29.7 cm sheet of paper which was fixed at the center of the patient’s sagittal midline. In the Letter Cancellation task, 60 target letters ‘A’ are distributed among other distractor letters. The Bells test requires identifying 35 bell icons distributed all over the sheet between other symbols. In these two cancellation tasks, patients were asked to cancel all of the targets, ‘A’ letters or bells respectively. The maximum duration of each test was not fixed in advance but depended on the patient being satisfied with his performance and confirming this twice. For the Letter and Bells Cancellation tasks, we calculated the Center of Cancellation (CoC) using the procedure by Rorden and Karnath (2010). The CoC is a sensitive measure capturing both the number of omissions, as well their location. For the lesion-behavior mapping, we calculated the mean CoC from the two cancellation tasks for each patient and used this score for our analyses. Visual field defects were examined by the common neurological confrontation technique.

### 2.3. Imaging

Structural imaging was acquired either by MRI (n = 106) or CT (n = 97), performed on average 3.5 days (SD = 4.6) after stroke-onset. If both imaging modalities were available, MR scans were preferred. In participants where MR scans were available, we used diffusion-weighted imaging (DWI) if the images were acquired within 48 h after stroke onset or T2-weighted fluid attenuated inversion recovery (FLAIR) images for later scans. Lesion boundaries were manually marked on the transversal slices of the individual MR or CT scans using the free MRIcron software (www.mccauslandcenter.sc.edu/mricro/mricron).

Normalization of CT or MR scans to MNI space with 1×1×1 mm resolution was performed by using the Clinical Toolbox (Rorden et al., 2012) under SPM8 (www.fil.ion.ucl.ac.uk/spm), and by registering lesions to its age-specific templates oriented in MNI space for both CT and MR scans (Rorden et al., 2012). If available, the MR scans were coregistered with a high resolution T1-weighted structural scan in the normalization process. Delineation of lesion borders and quality of normalization were verified by consensus of always two experienced investigators (one of them H.-O.K.). An overlap of all normalized lesions is shown in Fig. 1. The average lesion size in the sample was 45.52 cm^3^ (SD = 52.67 cm^3^). In the supplemental material we show overlap plots of normalized lesions separated for each imaging modality (Fig. S2) as well as a histogram of the lesion size distribution (Fig. S3 B).

**Figure 1:**
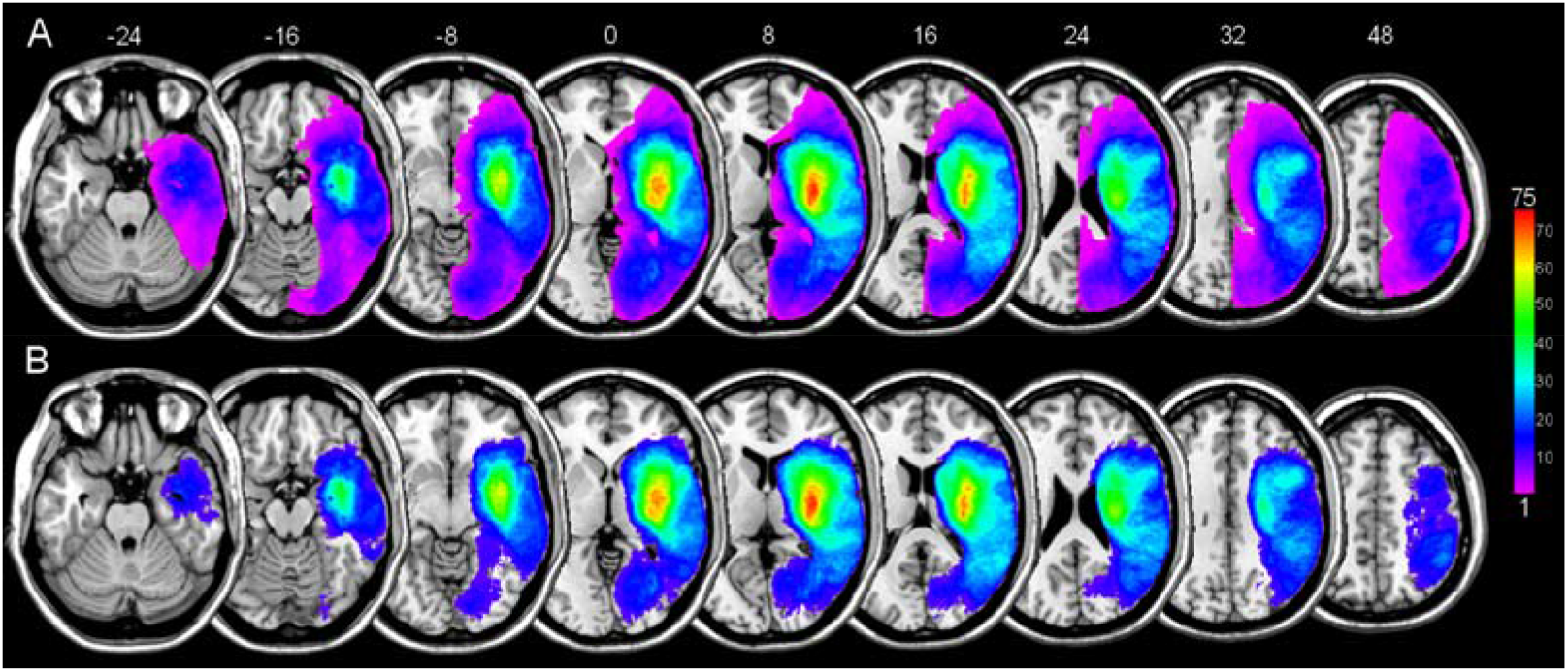
Topography of brain lesions. **A:** Simple lesion overlap topography of all 203 patients. **B:** Lesion overlap topography showing only voxels within the voxel mask for statistical testing with at least 10 patients having a lesion. The colorbar indicates the number of overlapping lesions (the peak of N = 75 represents 37% of the total sample). Numbers above the slices indicate z-coordinates in MNI space.

### 2.4. Multivariate lesion-behavior mapping

#### 2.4.1. Support Vector Regression

For our analysis, we implemented a multivariate lesion-symptom mapping method based on support vector regression (SVR) (Drucker et al., 1996; Vapnik, 1995). Lesion mapping based on support vector regression employs supervised machine learning algorithms to develop a model based on training input data which best describes the continuous relationship between behavioral scores and lesion location. Hence, it can be seen as an extension of Support Vector Machines (SVM) (Cortes and Vapnik, 1995) used for classifying data sets into different categories. Support vector regression based lesion-symptom mapping (SVR-LSM) has already been implemented and validated in a synthetic dataset, in a real dataset composed of aphasic patients (Zhang et al., 2014), as well as two recent publications (DeMarco and Turkeltaub, 2018; Sperber et al., in press; Chen et al., 2018). Moreover, SVR-LSM can be employed on a whole-brain (voxel-wise) level, using the full range of lesion information while respecting the relationships between multiple voxels simultaneously and is thus able to provide high resolution lesion-symptom mapping topographies.

#### 2.4.2. Data Analysis

The analysis was performed with MATLAB 2016a and libSVM Vers. 3.21 (Chang and Lin, 2013). We used a publicly available collection of scripts (https://github.com/yongsheng-zhang/SVR-LSM) employed in the study by Zhang et al. (2014) and adopted algorithms for control for lesion size and for the derivation of a topography from SVR β-parameters. For the detailed methodological procedure and theoretical background of SVR-LSM in general, see Zhang et al. (2014). Only voxels where at least 10 patients had a lesion were included in the analysis and constituted the voxel mask for statistical testing. Exclusion of voxels with infrequent lesion affection was performed to restrict the analysis to voxels with reasonable statistical power and thus to prevent that results are biased by brain regions that are only rarely affected (Karnath et al., 2018). The employed analysis is therefore no strict whole-brain analysis, but – contrary to region-of-interest analyses – it allows an unbiased investigation of all brain areas that contain a certain degree of information. First, the lesion status of each participant was regrouped into a column vector. To control for lesion size, each vector was then normalized to have a unit norm, a procedure also known as direct total lesion volume control (dTLVC) (Zhang et al., 2014). Lesion volume control is an important preprocessing step as the severity of a symptom is generally related to the lesion size, as shown for our data (r = 0.54; p < 0.001) in Fig. S3 A in the supplemental material. To estimate the SVR hyperplane and project our initial data into a higher dimensional space, we implemented an epsilon-SVR model and used a non-linear radial basis function (RBF) Kernel. In order to improve the performance of the learning algorithms and to choose a model best describing our data, a preselection of the model hyperparameters cost (C) and gamma (γ) needs to be done. Following general recommendations in the libSVM toolbox manual (Chang and Lin, 2013), we added an optimization procedure using grid search. The range of investigated parameters was chosen as in the study by Zhang et al., 2014: C = 1, 10, 20, 30, 40, 50, 60, 70, 80, and γ = 0.1, 1, 2, 3, 4, 5, 6, 7, 8, 9, 10, 15, 20, 25, 30. Using a five-fold cross-validation scheme, we evaluated both prediction accuracy and reproducibility of each parameter combination (see Zhang et al., 2014). During this procedure, the whole dataset is separated into 4/5 training data, which is used to generate the multivariate model. Then this model is tested on the unknown leftover 1/5 of the data to prevent overfitting and to get a good estimate of the performance of the model to unknown data. To save computational power, we reduced the number of iterations from 40 to 5 compared to the initial analysis in Zhang et al., 2014 and evaluated mean prediction accuracy and reproducibility scores, based on these 5 iterations for each parameter set. We define mean prediction accuracy, as in Zhang et al. (2014), to be the mean correlation coefficient between predicted scores and out of sample testing scores of 5 times 5-fold cross-validations. Note that for each of the 5 iterations, new random subsets of training and testing scores were drawn from the whole dataset. After SVR model construction, β-parameters are remapped onto a three-dimensional brain topography allowing us to derive the reproducibility score by calculation of the mean correlation coefficient between any two SVR-LSM β-parameter maps from the drawn subsets. Finally, using the best combination of C and γ for model construction, the remapped β-parameters are tested by using a permutation approach, comparing the SVR β-parameters voxel-wise with new β-parameters drawn for each permutation through randomization of behavioral scores. Results are reported with correction for multiple comparisons that survived a False Discovery Rate (Benjamini and Yekutieli, 2001) (FDR) correction at q = 0.05, determined by 10000 permutations. As statistical testing is performed on a voxel-by-voxel basis, a form of multiple comparison correction is required to prevent an increase of false alarms (Sperber et al., in press).

### 2.5. Univariate voxel-based lesion behavior mapping

To compare the SVR-LSM technique with traditional analyses, we also performed mass-univariate VLBM analyses on the same data set. As for MLBM, only voxels where at least 10 patients had a lesion were included in the analysis and constituted the voxel mask for statistical testing. The variants of lesion volume correction and correction for multiple comparisons differ between univariate studies. Nevertheless, the exact choice might have an impact on the topographical outcome of the univariate results as shown recently by Pustina and colleagues (2018). Therefore, we decided to show results using different parameter configurations, providing a small cross section of what is currently employed in the field. Hence, for the univariate analyses, there were, in total, 4 configurations: A) without correction for lesion size including family-wise error correction (FWE) for multiple comparisons based on 10000 permutations at p < 0.05; B) with correction for lesion size – by regressing lesion size out of behavior – including FWE correction for multiple comparisons based on 10000 permutations at p < 0.05; C) without correction for lesion size including False Discovery Rate (FDR) correction for multiple comparisons at q = 0.05; D) with correction for lesion size – by regressing lesion size out of behavior – including FDR correction for multiple comparisons at q = 0.05. All univariate analyses were carried out using the NiiStat tool (https://github.com/neurolabusc/NiiStat) and were based on the general linear model (identical to a Student’s pooled-variance t-test).

### 2.6. Supplementary analysis

While the need of lesion volume correction is widely accepted, the exact technique to be used is still under discussion. Zhang and colleagues (2014) validated the dTLVC method for real and synthetic lesion data and argued that a regression based correction might be excessively conservative. On the other hand, DeMarco and Turkeltaub (2018) put forward that the dTLVC method might be too liberal and advocated for using regression based correction. To address both positions and concerns, we implemented a supplemental analysis using the same parameters as for our main SVR-LSM analysis, except for now regressing lesion size out of both behavioral and lesion scores instead of the dTLVC procedure. Scripts for this supplemental analysis have been adopted from a recently published toolbox and are available online (github.com/atdemarco/svrlsmgui; DeMarco and Turkeltaub, 2018).

### 2.7. Atlas Overlap

Labeling of all the resulting voxel-wise statistical maps with respect to grey matter brain regions was done by overlaying the maps on the Automatic Anatomical Labelling atlas (AAL; Tzourio-Mazoyer et al., 2002) distributed with MRIcron. The localization of white matter fiber tracts damaged by the lesion was based on two different fiber tract atlases: the Juelich probabilistic cytoarchitectonic atlas (Bürgel et al., 2006) as well as the tractography-based probabilistic fiber atlas (Thiebaut De Schotten et al., 2011). We decided to interpret the data according to these two WM atlases simultaneously due to the marked variance between DTI- and histology-based white matter atlases (de Haan and Karnath, 2017). The WM probabilistic maps were thresholded at p>= 0.3 before being overlaid on the statistical topography.

## 3. Results

### 3.1. Parameter optimization

Testing different sets of C and γ combinations, we got a similar pattern of that what has already been reported in previous investigations (Rasmussen et al., 2012; Zhang et al., 2014). Thus, the C and γ variables provide a trade-off between prediction accuracy and reproducibility (Fig. 2 A and B). Without having any reference about a certain standard in choosing the right parameter combination, we decided to perform the final analysis with a C of 30 and γ of 4, as these values provided, compared to the other combinations, a decent prediction accuracy (0.43) while keeping the reproducibility index (0.91) as high as possible. With this combination, the model achieves an R^2^ of 0.19.

**Figure 2:**
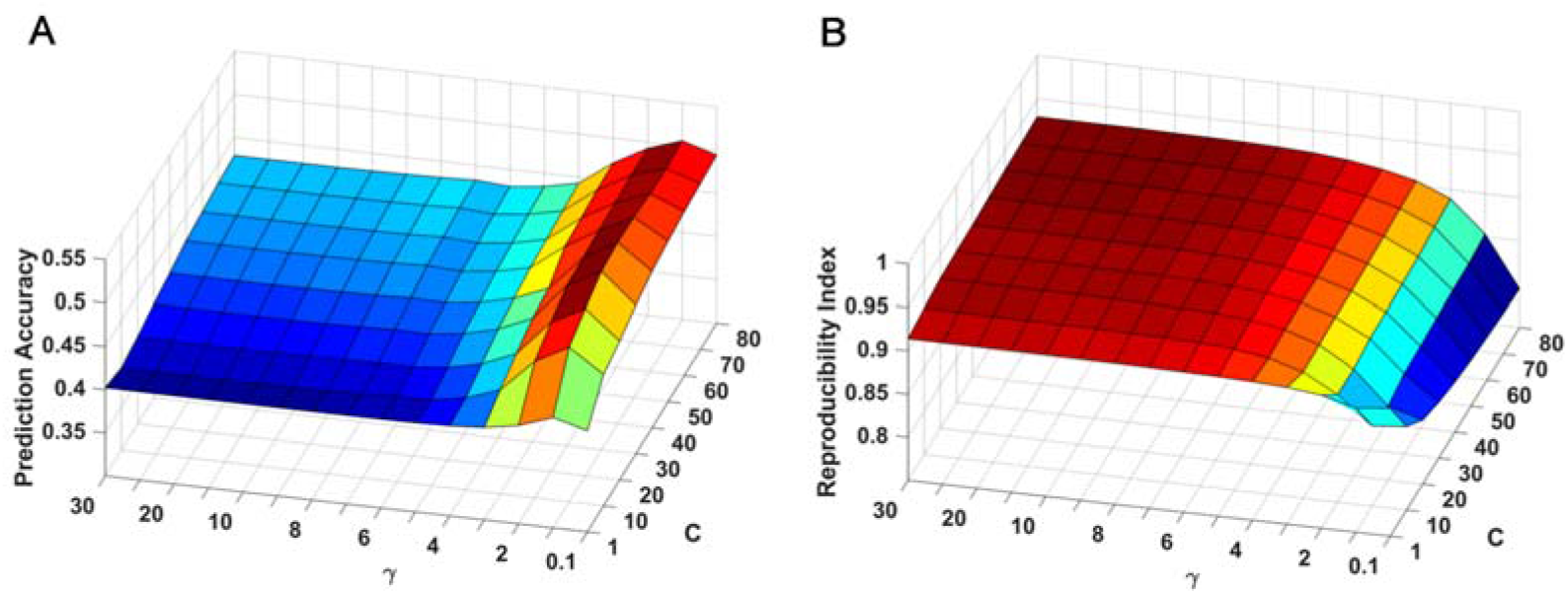
Estimation of best hyper-parameters C and γ. SVR-LSM parameter estimation results. Prediction Accuracy (**A**) and Reproducibility Index (**B**) (see Rasmussen et al., 2012; Zhang et al., 2014) are plotted for the different sets of C and γ parameters to find the optimal combination.

### 3.2. Multivariate lesion-behavior relationships

Resulting topographies of the SVR-LSM analysis using continuous CoC scores revealed a perisylvian network including parietal, frontal and temporal grey matter regions as well as interconnecting WM fibers (Fig. 3). An exact overview of the grey and white matter structures significantly involved and showing at least 100 mm3 overlapping voxels with the respective atlas structures is given in Table 2. Large clusters incorporated middle and superior temporal gyri as well as the inferior parietal lobule, including angular and supramarginal gyri. Smaller clusters affected inferior and middle frontal gyri, as well as the pre- and postcentral gyri. Moreover, significant lesion patterns included the insula and subcortical structures such as the pallidum, putamen and caudate nucleus. The overlap with both WM atlases consistently showed significant clusters affecting the uncinate fasciculus and the inferior occipito-frontal fasciculus. In addition, only the WM atlas by Thiebaut De Schotten et al. (2011) implicated the inferior longitudinal, as well as the superior longitudinal/arcuate fasciculus and the internal capsule, while only the Juelich WM atlas identified the superior occipito-frontal fasciculus. In addition, Fig. 3D shows the thresholded β–map of the SVR-LSM analysis. Please note, that individual β-values should be interpreted with caution as they don’t provide a one to one inference to anatomical localization and cannot be considered as classical test statistics (see also Sperber et al., in press), as for example t- or z-values. In general, β-values are correlating with probability measures (e.g. high β results in low p-value). However, this is not true for all tested voxels and it is thus possible that a low β becomes significant or vice-versa, a relatively high β might not reach statistical significance. An illustration of this can be found in Fig. S4.

**Figure 3:**
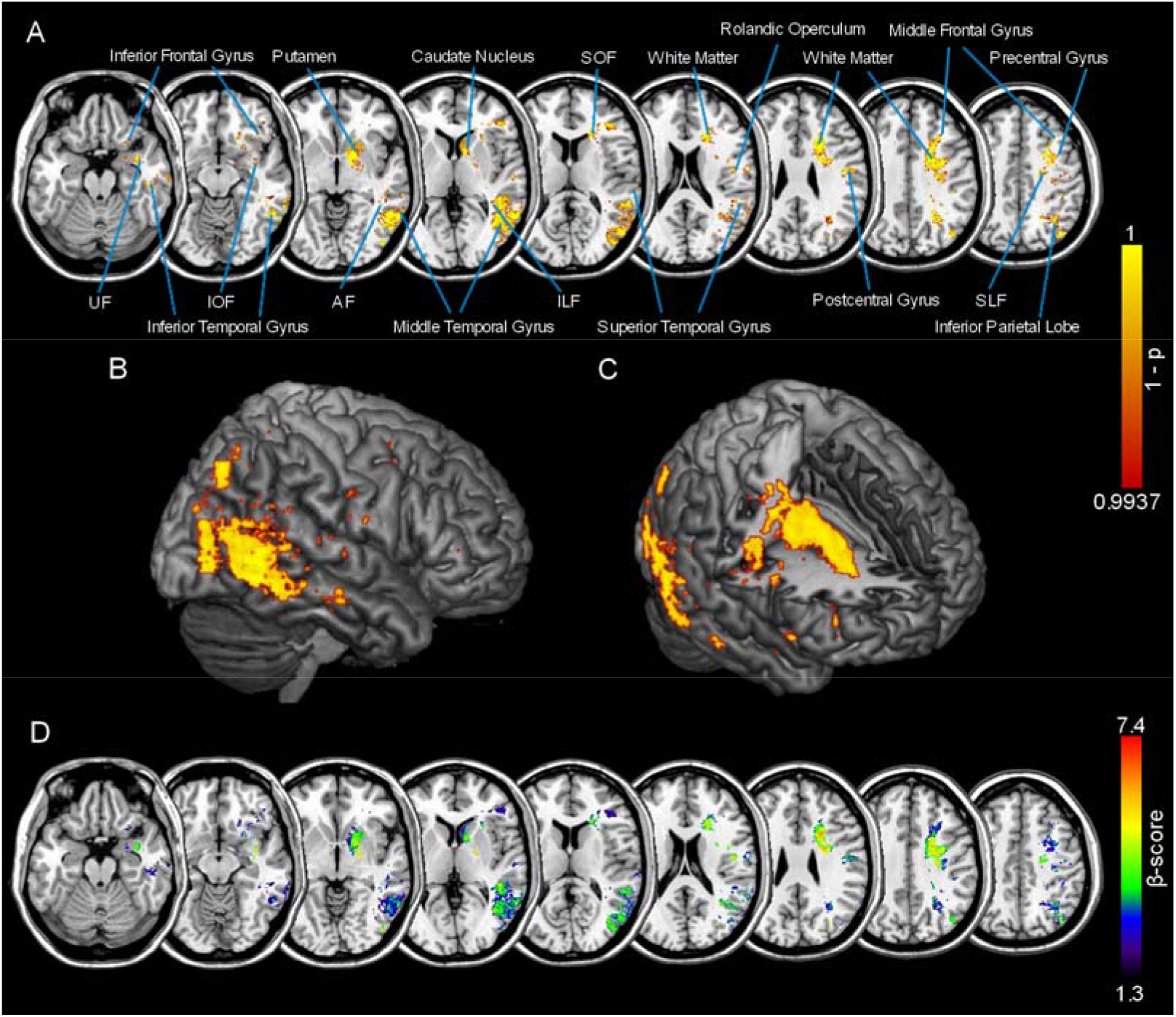
Results of the multivariate lesion-behavior mapping. Support vector regression based multivariate lesion-symptom mapping results using data of 203 patients. Lesion volume correction was performed by applying dTLVC. **A:** Permutation-thresholded statistical map of SVR-LSM on CoC scores (FDR-corrected at q = 0.05, corresponding to a threshold of p < 0.0063), illustrating the anatomical regions significantly associated with the core deficit of spatial neglect. Significant clusters were interpreted according to the AAL atlas (Tzourio-Mazoyer et al., 2002) for grey matter regions and to the Juelich probabilistic cytoarchitectonic fiber tract atlas (Bürgel et al., 2006) as well as the tractography-based probabilistic fiber atlas by Thiebaut De Schotten et al. (2011) for white matter structures. **B and C:** three-dimensional renderings of the same map using the 3D-interpolation algorithm provided by MRIcron (http://people.cas.sc.edu/rorden/mricron/index.html; 8mm search depth) with sagittal view for **B** and inside view for **C**. Results of **A, B** and **C** are shown as 1-p. **D:** Thresholded β-parameter map showing only significant areas according to **A.** Abbreviations: SLF – superior longitudinal fasciculus; AF – arcuate fasciculus; ILF – inferior longitudinal fasciculus; IOF – inferior occipitofrontal fasciculus; SOF – superior occipitofrontal fasciculus; UF – Uncinate fasciculus.

**Table 2:** Detailed overview of all significant grey and white matter clusters – MLBM. Overlap of MLBM analysis with grey and white matter atlases (FDR-corrected at q = 0.05, corresponding to a threshold of p < 0.0063). For grey matter structures, reports are generated using the Automatic Anatomical Labeling atlas (AAL; Tzourio-Mazoyer et al., 2002). For white matter structures, reports are generated using the Juelich probabilistic cytoarchitectonic atlas (Bürgel et al., 2006) and the tractography-based probabilistic fiber atlas by Thiebaut De Schotten et al. (2011) defined at a probability of p >= 0.3. Only structures with at least 100 mm3 of overlapping voxels were reported in the table.

| GM structure (AAL) | Number of voxels (mm<sup>3</sup>) | $\beta$ (Mean/SD) | Peak $\beta$ |
| --- | --- | --- | --- |
| Middle temporal gyrus | 8441 | 3.13/0.60 | 5.66 |
| Inf. temporal gyrus | 3078 | 2.59/0.57 | 4.40 |
| Middle occipital gyrus | 1677 | 3.37/0.61 | 5.73 |
| Angular gyrus | 1398 | 3.14/0.58 | 4.45 |
| Sup. temporal gyrus | 968 | 3.01/0.54 | 5.10 |
| Pallidum | 848 | 4.35/1.09 | 7.03 |
| Middle frontal gyrus | 739 | 2.62/0.53 | 4.73 |
| Caudatum | 719 | 3.24/0.63 | 5.72 |
| Inf. frontal gyrus/triangular | 706 | 2.23/0.23 | 3.99 |
| Postcentral gyrus | 570 | 2.92/0.35 | 4.17 |
| Putamen | 481 | 3.71/0.74 | 6.54 |
| Rolandic operculum | 422 | 3.74/0.53 | 5.22 |
| Precentral gyrus | 346 | 3.07/0.66 | 5.38 |
| Inf. frontal gyrus/orbital | 314 | 3.80/0.58 | 4.41 |
| Amygdala | 279 | 2.96/0.49 | 5.25 |
| Supramarginal gyrus | 228 | 2.88/0.35 | 3.73 |
| Inf. occipital gyrus | 214 | 3.45/0.95 | 5.23 |
| Inf. parietal gyrus | 166 | 2.81/0.24 | 3.68 |
| Insula | 119 | 3.00/0.93 | 5.99 |
| Hippocampus | 115 | 3.07/0.64 | 4.87 |
| Sup. temporal pole | 113 | 2.54/0.54 | 3.98 |

| WM structure (Juelich) | Number of voxels (mm<sup>3</sup>) | $\beta$ (Mean/SD) | Peak $\beta$ |
| --- | --- | --- | --- |
| Callosal body | 2703 | 4.10/1.09 | 7.31 |
| Optic radiation | 1393 | 3.07/0.67 | 5.48 |
| Corticospinal tract | 832 | 3.52/0.98 | 6.14 |
| Sup. occipito-frontal fasciculus | 262 | 4.26/0.91 | 6.84 |
| Inf. occipito-frontal fasciculus | 168 | 4.56/0.56 | 6.30 |
| Uncinate fasciculus | 134 | 4.36/0.68 | 6.30 |

| WM structure (Thiebaut de Schotten) | Number of voxels (mm<sup>3</sup>) | $\beta$ (Mean/SD) | Peak $\beta$ |
| --- | --- | --- | --- |
| Corpus callosum | 3245 | 4.05/1.01 | 7.39 |
| Internal capsule | 2846 | 4.28/1.05 | 7.23 |
| Inf. longitudinal fasciculus | 2545 | 3.10/0.67 | 5.48 |
| Arcuate fasciculus | 2315 | 3.15/0.61 | 5.42 |
| Posterior segment (Arcuate) | 2036 | 3.07/0.58 | 4.88 |
| Corticospinal tract | 1845 | 4.19/1.07 | 7.23 |
| Uncinate fasciculus | 1249 | 3.26/0.82 | 6.30 |
| Cingulum | 1200 | 3.39/0.61 | 5.37 |
| Anterior commissure | 1177 | 3.21/0.85 | 5.88 |
| Inf. occipito-frontal fasciculus | 798 | 3.49/0.81 | 5.98 |
| Optic radiations | 306 | 3.37/0.66 | 5.42 |
| Anterior segment (Arcuate) | 282 | 3.71/0.57 | 5.42 |
| Sup. longitudinal fasciculus | 144 | 3.04/0.33 | 4.87 |

### 3.3. Univariate voxel-based lesion-behavior relationships

The VLBM analysis including family-wise error correction (FWE) for multiple comparisons without correction for lesion size (Fig. 4A) revealed mainly involvement of inferior and middle frontal as well as middle and superior temporal cortical grey matter areas. Moreover, affection of subcortically the putamen, caudate and pallidum can be delineated; parietal cortex was not involved. A detailed overview of grey and white matter structures significantly involved and with at least 100 mm^3^ overlap with the respective atlases is given in Table 3A. The comparison with white matter atlases reveals involvement of the superior occipito-frontal fasciculus for the Juelich WM atlas as well as the arcuate and uncinate fasciculi for the WM atlas by Thiebaut De Schotten et al. (2011). Moreover, the latter delineated affection of the inferior occipito-frontal and longitudinal fasciculi, as well as the internal capsule.

**Figure 4:**
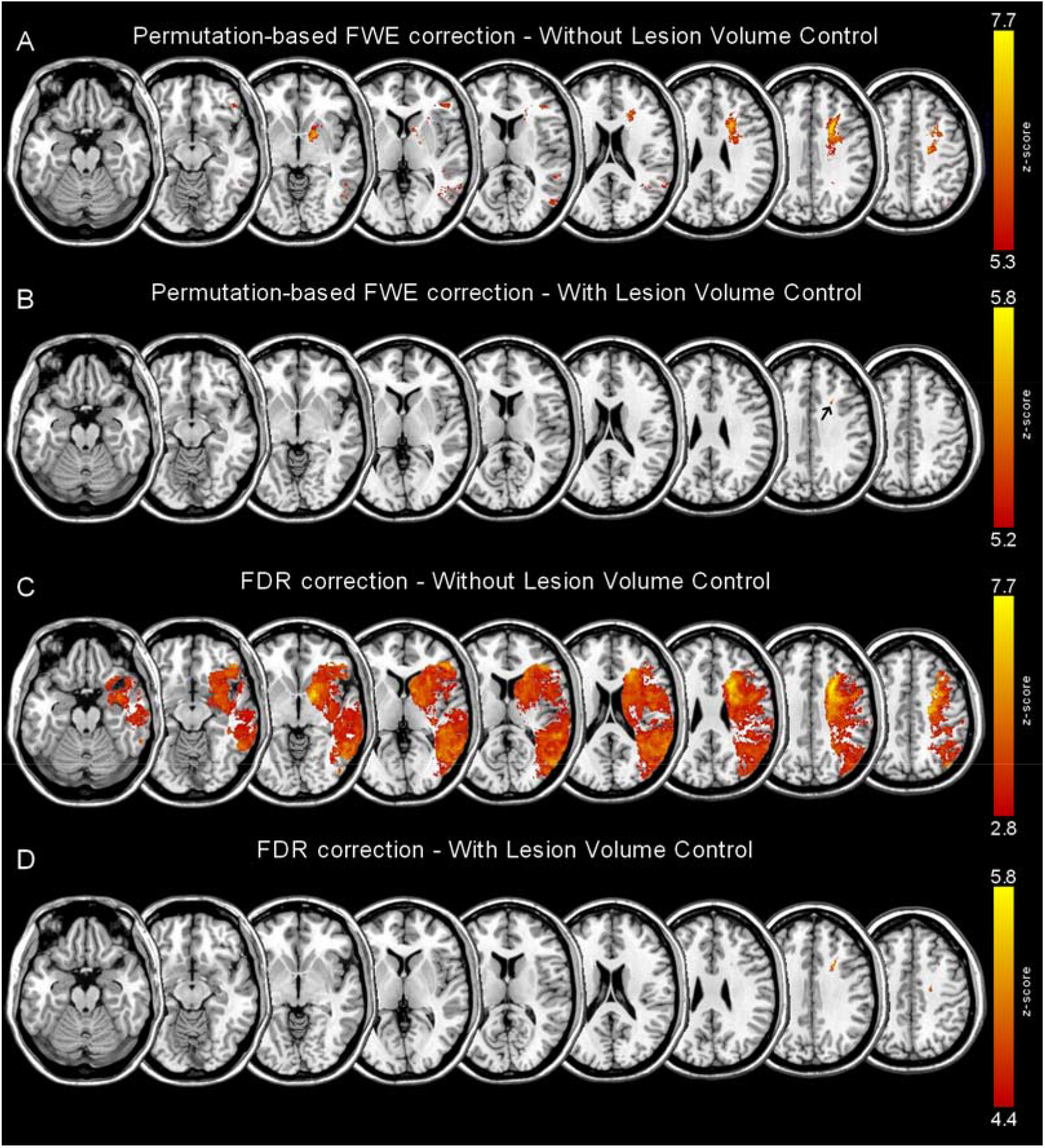
Results of the univariate lesion-behavior mapping. Mass-univariate lesion-symptom mapping results using data of 203 patients. Z-score maps are plotted for FWE permutation-thresholded as well as FDR-thresholded VLBM analyses with and without lesion volume correction on CoC scores. **A:** FWE permutation thresholded VLBM analysis without correction for lesion size at p < 0.05, corresponding to a threshold of z > 5.3475; **B:** FWE permutation thresholded VLBM analysis with correction for lesion size – by regressing lesion size out of behavior – at p < 0.05, corresponding to a threshold of z > 5.2251; **C:** FDR thresholded VLBM analysis without correction for lesion size at q = 0.05, corresponding to a threshold of z > 2.8607; **D:** FDR thresholded VLBM analysis with correction for lesion size – by regressing lesion size out of behavior – at q = 0.05, corresponding to a threshold of z > 4.4772.

**Table 3:** Detailed overview of all significant grey and white matter clusters – VLBM. Overlap of VLBM analysis without control for lesion size with grey and white matter atlases. **A:** permutation-based FWE correction at p < 0.05, corresponding to a threshold of z > 5.3475; **B:** FDR correction at q = 0.05, corresponding to a threshold of z > 2.8607. For grey matter structures, reports are generated using the Automatic Anatomical Labeling atlas (AAL; Tzourio-Mazoyer et al., 2002). For white matter structures, reports are generated using the Juelich probabilistic cytoarchitectonic atlas (Bürgel et al., 2006) and the tractography-based probabilistic fiber atlas by Thiebaut De Schotten et al. (2011) defined at a probability of p >= 0.3. Only structures with at least 100 mm3 of overlapping voxels were reported in the table. For the VLBM analyses with correction for lesion size, no overlap table is generated, as no clear labeling was possible.

| **A) FWE- thresholding without correction for lesion size** |  |  |  |
| --- | --- | --- | --- |
| GM structure (AAL) | Number of voxels (mm<sup>3</sup>) | z-score (Mean/SD) | Peak z-score |
| Middle temporal gyrus | 1304 | 5.66/0.28 | 6.86 |
| Pallidum | 523 | 5.93/0.48 | 7.26 |
| Middle frontal gyrus | 438 | 5.85/0.35 | 6.96 |
| Inf. frontal gyrus/triangular | 391 | 5.63/0.19 | 6.05 |
| Putamen | 228 | 5.66/0.33 | 6.94 |
| Sup. temporal gyrus | 167 | 5.58/0.20 | 6.23 |
| Caudatum | 129 | 5.57/0.19 | 6.20 |
| Inf. frontal gyrus/orbital | 113 | 5.60/0.20 | 6.06 |
| Middle occipital gyrus | 113 | 5.73/0.28 | 6.55 |
| WM structure (Juelich) | Number of voxels (mm<sup>3</sup>) | z-score (Mean/SD) | Peak z-score |
| Callosal body | 1832 | 6.07/0.53 | 7.66 |
| Corticospinal tract | 544 | 5.80/0.36 | 6.75 |
| Sup. occipito-frontal fasciculus | 128 | 5.78/0.37 | 6.90 |

| WM structure<br>(Thiebaut De Schotten) | Number of voxels (mm<sup>3</sup>) | z-score (Mean/SD) | Peak z-score |
| --- | --- | --- | --- |
| Corpus callosum | 2508 | 6.01/0.52 | 7.66 |
| Internal capsule | 2187 | 5.97/0.49 | 7.66 |
| Corticospinal tract | 1771 | 5.89/0.41 | 7.41 |
| Cingulate | 530 | 5.78/0.34 | 7.24 |
| Anterior commissure | 304 | 5.97/0.48 | 7.26 |
| Arcuate fasciculus | 269 | 5.63/ 0.28 | 6.58 |
| Posterior segment (Arcuate) | 267 | 5.63/ 0.28 | 6.58 |
| Cortico ponto cerebellum | 169 | 5.76/ 0.34 | 6.69 |
| Uncinate | 126 | 5.60/ 0.22 | 6.36 |
| Inf. longitudinal fasciculus | 122 | 5.63/ 0.29 | 6.68 |
| Inf. occipito-frontal fasciculus | 100 | 5.58/ 0.22 | 6.36 |

**B) FDR- thresholding without correction for lesion size**
| GM structure (AAL) | Number of voxels (mm<sup>3</sup>) | z-score (Mean/SD) | Peak z-score |
| --- | --- | --- | --- |
| Middle temporal gyrus | 25761 | 4.03/0.73 | 6.86 |
| Sup. temporal gyrus | 14260 | 3.68/0.61 | 6.23 |
| Angular gyrus | 8455 | 3.65/0.52 | 5.86 |
| Putamen | 7628 | 4.02/0.65 | 6.94 |
| Insula | 7501 | 3.45/0.40 | 5.36 |
| Middle occipital gyrus | 6529 | 3.69/0.58 | 6.55 |
| Inf. temporal gyrus | 6374 | 4.00/0.72 | 6.72 |
| Inf. frontal gyrus/triangular | 5894 | 3.89/0.77 | 6.05 |
| Caudatum | 5335 | 4.18/0.59 | 6.20 |
| Precentral gyrus | 5335 | 3.72/0.64 | 6.85 |
| Inf. frontal gyrus/opercular | 5158 | 3.47/0.46 | 5.87 |
| Middle frontal gyrus | 5078 | 4.08/0.81 | 6.96 |
| Rolandic operculum | 4609 | 3.60/0.54 | 5.90 |
| Inf. frontal gyrus/orbital | 4433 | 4.02/0.58 | 6.06 |
| Supramarginal gyrus | 4196 | 3.36/0.40 | 5.68 |
| Inf. parietal gyrus | 2814 | 3.35/0.38 | 4.89 |
| Postcentral gyrus | 2776 | 3.31/0.35 | 5.19 |
| Pallidum | 2158 | 4.87/0.80 | 7.26 |
| Sup. temporal pole | 2123 | 3.42/0.47 | 5.61 |
| Amygdala | 1351 | 3.96/0.59 | 6.12 |
| Hippocampus | 1060 | 3.57/0.58 | 5.60 |
| Sup. parietal gyrus | 704 | 3.25/0.30 | 4.72 |
| Thalamus | 664 | 3.42/0.47 | 5.14 |
| Middle temporal pole | 601 | 3.54/0.53 | 5.47 |
| Inf. occipital gyrus | 392 | 4.12/0.83 | 6.50 |
| Olfactory cortex | 276 | 3.62/0.38 | 4.77 |
| Sup. occipital gyrus | 195 | 3.18/0.26 | 4.21 |
| Fusiform gyrus | 154 | 3.40/0.45 | 4.81 |
| Transverse temporal gyrus | 139 | 3.03/0.13 | 3.48 |
| Sup. frontal gyrus/orbital | 103 | 3.94/0.43 | 4.86 |

| WM structure (Juelich) | Number of voxels (mm<sup>3</sup>) | z-score (Mean/SD) | Peak z-score |
| --- | --- | --- | --- |
| Corticospinal tract | 10278 | 3.90/0.76 | 6.75 |
| Corpus callosum | 8367 | 4.40/1.10 | 7.66 |
| Optic radiation | 7702 | 3.79/0.59 | 6.64 |
| Sup. longitudinal fasciculus | 1925 | 3.64/0.53 | 6.11 |
| Inf. occipito-frontal fasciculus | 1742 | 3.72/0.49 | 5.14 |
| Sup. occipito-frontal fasciculus | 1377 | 4.35/0.75 | 6.90 |
| Acoustic radiation | 794 | 3.32/0.36 | 4.68 |
| Uncinate fasciculus | 738 | 3.98/0.46 | 5.80 |

| WM structure<br>(Thiebaut De Schotten) | Number of voxels (mm<sup>3</sup>) | z-score (Mean/SD) | Peak z-score |
| --- | --- | --- | --- |
| Internal capsule | 16030 | 4.24/0.95 | 7.66 |
| Arcuate fasciculus | 15722 | 3.91/0.63 | 6.58 |
| Corticospinal tract | 11583 | 4.23/0.97 | 7.41 |
| Inf. longitudinal fasciculus | 10365 | 3.80/0.60 | 6.68 |
| Posterior segment (Arcuate) | 9743 | 4.07/0.65 | 6.58 |
| Corpus callosum | 9737 | 4.59/1.06 | 7.66 |
| Inf. occipito-frontal fasciculus | 9049 | 3.84/0.58 | 6.36 |
| Uncinate fasciculus | 6650 | 3.97/0.61 | 6.36 |
| Anterior segment (Arcuate) | 6335 | 3.63/0.48 | 5.47 |
| Anterior commissure | 3889 | 4.17/0.79 | 7.26 |
| Optic radiations | 2965 | 3.84/0.58 | 5.79 |
| Cingulum | 2914 | 4.38/0.92 | 7.24 |
| Cortico ponto cerebellar | 2607 | 3.78/0.80 | 6.69 |
| Fornix | 1208 | 3.47/0.44 | 5.18 |
| Sup. longitudinal fasciculus | 1205 | 3.78/0.52 | 5.35 |
| Long segment (Arcuate) | 691 | 3.58/0.44 | 5.02 |

The VLBM analysis including family-wise error correction (FWE) and a correction for lesion size (Fig. 4B) revealed only a small cluster within frontal white matter. The univariate analysis using FDR-thresholding without correction for lesion size is showing a wide spreading map with 56% of the tested voxels becoming significant, spanning over frontal, temporal, parietal and occipital as well as subcortical areas and interconnecting white matter fibres (Fig. 4C). A detailed overview of grey and white matter structures significantly involved and with at least 100 mm3 overlap with the respective atlases is given in Table 3B. On the other side, employing lesion volume correction, the analysis with FDR-thresholding showed a rather conservative pattern – similar to FWE-thresholding with correction for lesion size – of two small clusters in the frontal white matter with no corresponding atlas label (Fig. 4D).

### 3.4. Supplementary analysis

The optimization procedure delineated a C of 1 and γ of 0.1 as optimal parameters. Results for the supplemental analysis revealed a much more conservative pattern as compared to the SVR-LSM analysis with dTLVC correction. If we regress lesion size out of both behavioral and lesion scores, the resulting topographies centered on several smaller nodes (see Supplementary Material; Fig. S1). One cluster of lesion symptom associations was found within the right basal ganglia (putamen, pallidum, head of caudate nucleus). Moreover, an anterior cluster was revealed within the white matter adjacent to inferior and middle frontal gyri. A further small node was found at the right middle/inferior temporal cortex.

## 4. Discussion

The present study examined the lesion-behavior relationship of spatial neglect in a newly recruited sample of right brain damaged patients. By employing an MLBM approach, we were able to delineate the postulated anatomy of the putative perisylvian network in one single analysis. In particular, it included the superior and middle temporal gyri, inferior parietal lobule, insula, and the inferior and middle frontal gyri. Subcortically, we observed affection of the pallidum, putamen and caudate nucleus as well as white matter fiber tracts, such as the superior and inferior longitudinal fasciculi, the superior and inferior occipito-frontal fasciculi, and the uncinate fasciculus.

To compare the results of the multivariate SVR-LSM technique with the traditional mass-univariate analysis technique, we conducted VLBM using different variants of lesion size correction and multiple comparisons correction, corresponding to the majority of the current univariate lesion-mapping studies. Despite the large sample size, the two univariate analyses with lesion volume control only detected one (FWE), respectively two (FDR) cluster(s) in frontal white matter. Only when keeping the residual signal of lesion size in the model, the univariate VLBM had enough power to generate a statistical map which was qualitatively comparable to the multivariate SVR-LSM map. In the VLBM analysis including FWE correction, the univariate processing detected most of the regions that were also detected by the SVR-LSM analysis, with the exception of the inferior parietal lobule. The major difference was that VLBM detected massively less signal, resulting in only few actually interpretable clusters. In contrast, the VLBM analysis including FDR thresholding showed a more liberal pattern. It was able to detect the regions discussed in the literature with the caveat that they were hidden within a considerable amount of probably false positive detections, limiting interpretation. It should be noted that the results of the latter two analyses were coupled to the omission of lesion volume control. However, since the severity of a behavioral symptom is often strongly correlated with total lesion volume and larger lesions are more likely to affect critical anatomical areas (Karnath et al., 2004), a form of correction is desired to detect the neural correlate specific to the behavioral symptom of interest. A further positive effect of lesion volume correction in VLBM analyses is that it reduces the systematic misplacement of the outcome (cf. Sperber and Karnath, 2017).

The simulations by Zhang and colleagues (2014) showed that SVR-LSM is characterized by a good receiver operator characteristic (ROC) performance, especially if lesion-volume correction is included. In contrast, ROC characteristics of VLBM were generally worse and thresholds with both good sensitivity and specificity were not available. This limits VLBM to a choice in favor of either sensitivity with accepting a larger number of false negatives (i.e. selecting a higher cutoff) or specificity with accepting a larger number of false positives (i.e. selecting a lower cutoff). Hence, our data might drive speculations about VLBM not being sensitive enough after permutation based FWE thresholding, or not specific enough after FDR based thresholding. However, simulation studies are required here. Taking all of our univariate findings together, one might conclude that in the framework of multi-area based syndromes, VLBM might not in general fail to detect such networks, but it suffers from different limitations. These might have contributed to the heterogeneous results of previous lesion-symptom mapping investigations in spatial neglect using mass-univariate VLBM (Karnath et al., 2004, 2011; Committeri et al., 2007; Sarri et al., 2009; Chechlacz et al., 2010; Saj et al., 2012; Thiebaut De Schotten et al., 2014; Rousseaux et al., 2015) and DTI/white matter fiber analyses (Thiebaut De Schotten et al., 2005; Urbanski et al., 2008, 2011; Karnath et al., 2009; Shinoura et al., 2009; Ciaraffa et al., 2013; Thiebaut De Schotten et al., 2014; Umarova et al., 2014; Lunven et al., 2015; Vaessen et al., 2016; Carter et al., 2017). While using meta-analytic approaches combining various VLBM results, one might be able to find all critical parts of the presumed network. The present study shows that MLBM analyses have the potential to achieve this by a single investigation while respecting recommendations on correction factors.

The two previous multivariate examinations on spatial neglect (Smith et al., 2013; Toba et al., 2017) were able to uncover only parts of this network. Very likely this is due to the small sample size in one of them (Toba et al., 2017) and – most importantly – the very limited number of a priori defined brain regions per multivariate model in both studies. In contrast, the present multivariate SVR-LSM analysis utilized the full voxel-wise information and uncovered a larger set of cortical and subcortical areas, able to account for inconsistencies on the anatomical representation of spatial neglect. The finding highly corresponds to the proposed ‘perisylvian network’ (Karnath, 2009), consisting of superior/middle temporal, inferior parietal, and ventrolateral frontal cortices, and representing the anatomical basis for processes involved in spatial orienting and neglect.

The right temporal lobe has been delineated in previous lesion mapping studies in spatial neglect (Karnath et al., 2001, 2004, 2011; Committeri et al., 2007; Saj et al., 2012; Rousseaux et al., 2015). Smith et al. (2013) found the superior temporal gyrus (STG) as being the only structure which contained unique information for predicting spatial neglect. Accordingly, the STG seems to play an important role in multisensory integration, conveying information from both, the dorsal route of visual information processing, as well as polysensory inputs from the ventral perceptual stream (for review, see Karnath, 2001). Further evidence for the importance of superior/middle temporal areas for spatial neglect comes from recent animal models (Bogadhi et al., 2019). The authors detected a causal relationship between spatial neglect like symptoms and the superior temporal sulcus by direct and indirect ‘inactivation’ of that area in the monkey brain, underlining it’s crucial role in covert attentional processing. In the human brain, the posterior part of the STG at the intersection to the inferior parietal cortex, an area which is called ‘temporo-parietal junction’ (TPJ) (Chang et al., 2013; Kincade et al., 2005; Macaluso and Doricchi, 2013) together with the ventral frontal cortex (VFC) (Corbetta and Shulman, 2002; Snyder and Chatterjee, 2006) have been reported as target areas for attentional reorienting, target detection and vigilance (Corbetta and Shulman, 2011). Lesions in these cortical areas, together with white matter disconnection hindering the information transmission between them (see below), seem to form the basis for the core deficit (see introduction section above) observed in spatial neglect patients.

The superior longitudinal fasciculus connects the ventral frontal cortex to parietal structures via different sub-branches, the SLF I, SLF II and SLF III, identified in both humans, and monkeys (Schmahmann and Pandya, 2006, Thiebaut De Schotten et al., 2011). The SLF has repeatedly been discussed as being a crucial fronto-parietal pathway for processes of attentional orienting (Corbetta and Shulman, 2002; Bartolomeo et al., 2007). The ventral branch of the SLF (SLF III) specifically connects brain regions within the ventral attention network (VAN) (Rushworth et al. 2006; Bartolomeo et al., 2012; Thiebaut De Schotten et al., 2011) engaged in the propagation of information of stimulus identity and during the automatic capture of spatial attention by visual targets (Corbetta and Shulman, 2002). A further intrahemispheric tract partly overlapping with the SLF and confirmed in the present investigation is the arcuate fasciculus (AF). The AF is sometimes considered as an additional subcomponent of the SLF (Makris et al., 2005; Vernooij et al., 2007) and has been discussed in the transmission of information related to visuospatial performance (Chechlacz et al., 2014; Ciaraffa et al., 2013; Thiebaut De Schotten et al., 2014). This fiber bundle is composed of long and short anterior as well as short posterior fibers connecting specifically perisylvian frontal, parietal and temporal areas (Catani and Thiebaut De Schotten, 2008). Additionally, our results indicate the crucial involvement of the occipitofrontal fasciculus (IOF) – also termed as ‘inferior frontooccipital fasciculus’ (IFOF) –, which runs through the temporal lobe medial to the lower insula and connects areas of the frontal cortex with posterior temporal, inferior parietal, and occipital cortices (Nieuwenhuys et al. 1988; Catani et al. 2002; Bürgel et al. 2006; Forkel et al., 2014; Lawes et al., 2008). It has been suggested, that damage of this tract may hamper the transmission and/or the serial encoding of visual information in memory (Humphreys et al., 2015). This might explain deficits in target/distractor discriminative cancellation tasks, whereas no specific link between the IOF on line cancellation without distractor items has been reported (Urbanski et al., 2008). Our analysis depicted also the inferior longitudinal fasciculus (ILF), a further association tract connecting temporal to occipital areas (Catani et al., 2008), which has been linked previously to spatial neglect (Bird et al., 2006; Toba et al., 2018).

In addition to the cortical network, significant clusters were also observed subcortically for the right basal ganglia, including putamen, pallidum, and caudate nucleus. Based on previous work using perfusion imaging to monitor the remote effects of subcortical lesions (e.g. Hillis et al., 2002; Karnath et al., 2005), it is very likely that lesions of the basal ganglia lead to behavioral dysfunction by impairing the cortical network indirectly by malperfusion. However, there might also be a direct involvement of the basal ganglia. A recent simulation study by Parr and Friston (2018) aimed to formulate spatial neglect as a computational deficit. By setting up a model structure corresponding to the anatomy of dorsal and ventral attention networks, as well as their subcortical contributions, they demonstrated that basal ganglia lesions can directly produce neglect behavior in a saccadic cancellation task.

Summarizing our anatomical results, multivariate lesion behavior mapping was able to depict the network by only one single analysis and by using only the CoC score as behavioral proxy for spatial neglect and attention, whereas previous studies needed to employ either a meta-analytical (Checlacz et al., 2012; Molenberghs et al., 2012), multi-imaging/multi-method (Corbetta et al., 2015; Ramsey et al., 2016), or ROI approaches (Smith et al., 2013) to increase power or to disentangle the behavioral sub-functions and map them separately (Verdon et al., 2010; Vaessen et al., 2016; Toba et al., 2018) to come to comparable conclusions. This indicates that the behavioral proxy we measured here is indeed a core symptom of spatial neglect as it might evolve regardless where within this network a lesion produces focal dysfunction or remote deficits through disconnection.

Nevertheless, we want to emphasize that future studies might apply the same multivariate lesion analysis procedure used here for the core symptom of spatial neglect to uncover the exact neural underpinnings of the dissociating spatial and non-spatial behaviors. To disentangle the different components, these behavioral tasks in such studies should be as fine graded as possible to detect a specific cognitive function in isolation (for discussion, see Sperber and Karnath, 2018; Vuilleumier, 2013; Saj et al., 2012).

As expected, the regression technique suggested by DeMarco and Turkeltaub (2018) to control for lesion size turned out to be much more conservative than the dTLVC procedure (Zhang et al., 2014). A comparison of the resulting statistical map (Supplementary Material; Fig. S1) to the patient overlap plot (Fig. 1) revealed that especially those areas with the highest lesion frequency across our entire sample were spared out. The regression technique suggested by DeMarco and Turkeltaub (2018) limited the ‘searchlight’ rather to the border areas of the space of interest. Hence, the regression based control for lesion size appears to reduce the amount of false positives significantly, but at the cost of a higher rate of false rejections. Indeed, DeMarco and Turkeltaub (2018) noted in their discussion, that the cost of using this technique is a more conservative voxel-wise thresholding. The authors applied SVR-LSM in combination with their lesion volume regression approach on twenty real behavioral scores, but found significant results only in seven of these analyses. This further underline that lesion volume control, by regressing out lesion size from both behavioral and lesion scores, might be excessively conservative in many situations (including the present data). Results using this type of lesion size control thus should be interpreted with caution.

Albeit all the benefits of the SVR-LSM approach exposed here, it is also important to point to limitations. While we have recently provided a detailed discussion of such issues (Karnath et al., 2018), we would like to highlight that multivariate lesion-mapping techniques – as traditional univariate methods – do not identify brain areas that are structurally intact, but dysfunctional due to (temporary) hypoperfusion or diaschisis. Both approaches are based on structural imaging only and might thus be meaningfully complemented by other imaging techniques, such as perfusion CT or MR perfusion-weighted imaging. Multimodal imaging of brain damage where structural and functional information is combined might provide a more accurate picture of the full extent of the network underlying a disturbed cognitive function. Secondly, a recent investigation has revealed that multivariate lesion behavior mapping is susceptible to misplacement of statistical topographies along the brain’s vasculature to about the same extent as mass-univariate VLBM analyses (Sperber et al., in press). Thus, despite the high potential of multivariate lesion behavior mapping methods, it is possible that the final topography is influenced by the dependence of damage status between voxels. Together with the abovementioned perfusion and/or direct neural dysfunction effects, this might explain findings in subcortical areas, including e.g. putamen and pallidum.

## 5. Conclusion

The comparison between univariate and multivariate lesion analysis techniques revealed that the detected signal was either very conservative or very liberal for VLBM, demonstrating clear benefits of a multivariate approach if behavior is organized in large networks. While control for lesion size led to only reduced signal with VLBM, the SVR-LSM approach uncovered a complex network pattern in one single analysis. The implementation of multivariate analysis techniques thus has potential to explain inconsistencies and controversies emerging in the literature on the anatomical underpinnings of neurological syndromes. Hence, our findings underline the importance of a right perisylvian network in spatial attention and specifically in the emergence of the core symptoms of spatial neglect. Therefore, the present work is a step forward to improve our understanding of both multivariate lesion-behavior mapping in general as well as of complex brain networks involved in spatial attention and orientation behavior in particular. A remaining task for future studies is to investigate if the SVR-LSM approach employed here is indeed the most suitable multivariate analysis technique for studying the type of research question addressed in the present study. A comparison of different multivariate algorithms in multivariate lesion-behavior mapping is needed.

## 6. Acknowledgments

This work was supported by the Deutsche Forschungsgemeinschaft (KA 1258/23-1) and National Institutes of Health (P50DC014664). Daniel Wiesen was supported by the Luxembourg National Research Fund (FNR/11601161); Christoph Sperber by the Friedrich Naumann Foundation.

## Abbreviations

VLBM –: Voxel-based lesion behavior mapping;
MLBM –: Multivariate lesion behavior mapping;
SVR –: Support vector regression;
SVR-LSM –: Support vector regression based lesion-symptom mapping;
FDR –: false discovery rate;
FWE –: Family Wise Error;
dTLVC –: Direct Total Lesion Volume Control

