## supplemental for "Using machine learning-based lesion behavior mapping to identify anatomical networks of cognitive dysfunction: spatial neglect and attention"

### Supplementary Material

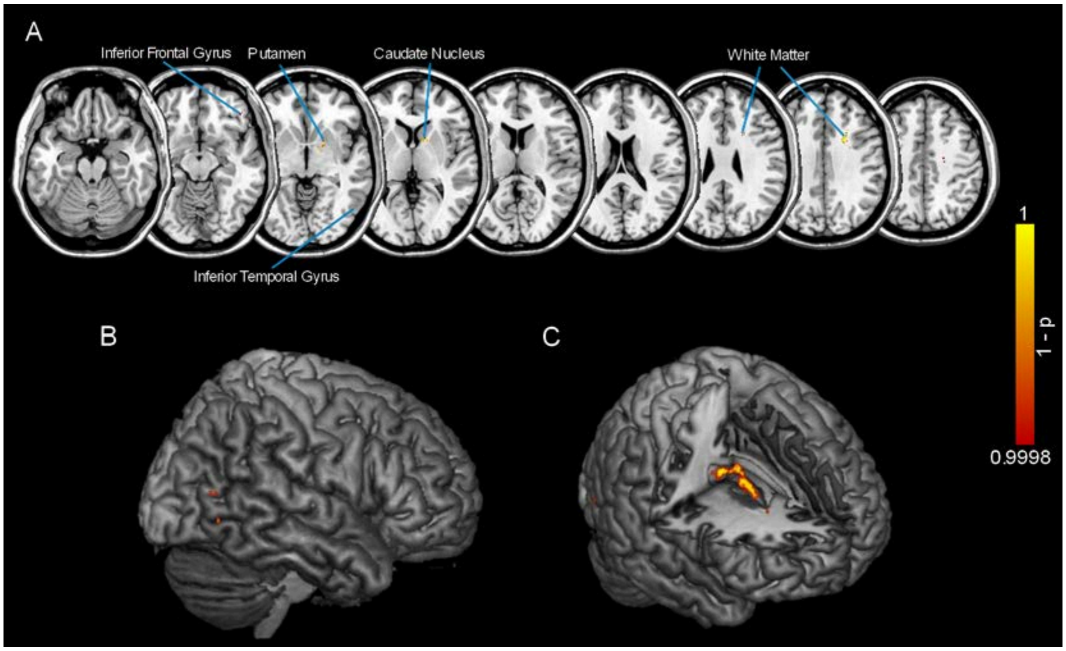


**Figure S1: Results of the multivariate lesion-behavior mapping controlled for lesion size by regression out of behavior and lesion**

Support vector regression based multivariate lesion-symptom mapping results using data of 203 patients. Lesion volume correction was performed by regressing lesion volume out of both behavioral and lesion scores (DeMarco and Turkeltaub, 2018). **A:** Permutation-thresholded statistical map of SVR-LSM on CoC scores (FDR-corrected at $q = 0.05$, corresponding to a threshold of $p < 0.0002$), illustrating the anatomical regions significantly associated with the core deficit of spatial neglect. Significant clusters were interpreted according to the AAL atlas (Tzourio-Mazoyer et al., 2002) for grey matter regions and to the Juelich probabilistic cytoarchitectonic fiber tract atlas (Bürgel et al., 2006) as well as the tractography-based probabilistic fiber atlas (Thiebaut De Schotten et al., 2011) for white matter structures. **B and C:**

three-dimensional renderings of the same map using the 3D-interpolation algorithm provided by MRIcron (http://people.cas.sc.edu/rorden/mricron/index.html; 8mm search depth) with sagittal view for **B** and **C**. Results are shown as 1-p.

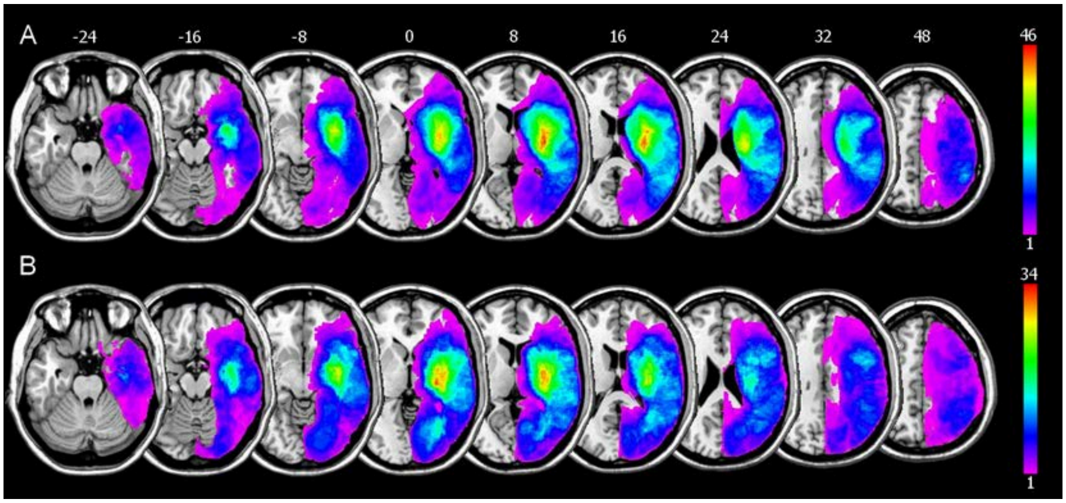


**Figure S2: Topography of brain lesions**

**A:** Lesion overlap topography of all lesions defined by MR (N = 106). **B:** Lesion overlap topography of all lesions defined by CT (N = 97). The colorbar indicates the number of overlapping lesions with peak at N = 46 for MR (i.e. 43% affection of all MR delineated lesions at peak) and N = 34 for CT (i.e. 35% affection of all CT delineated lesions at peak). Numbers above the slices indicate z-coordinates in MNI space.

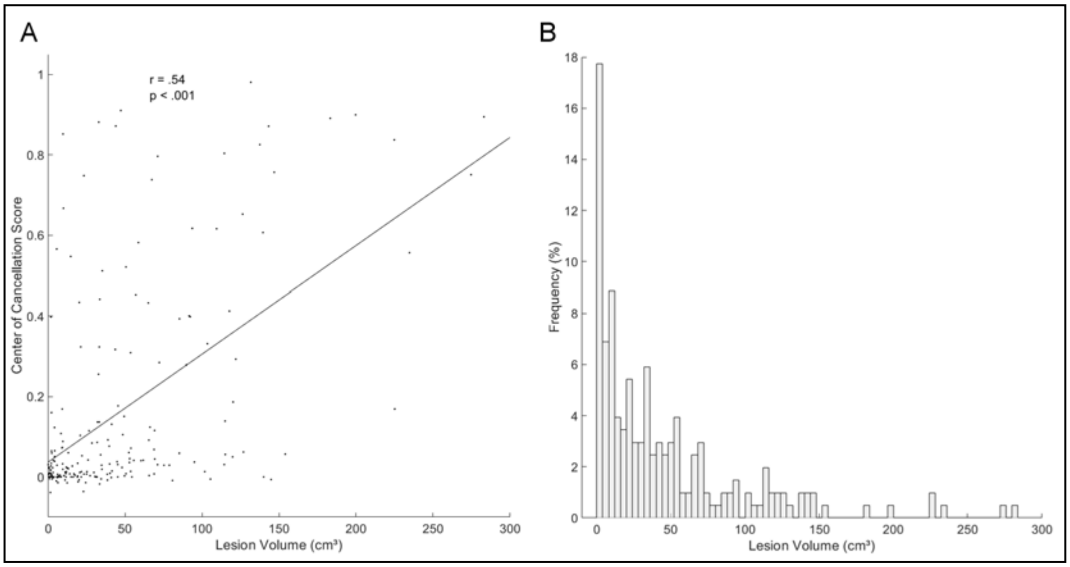


**Figure S3: Lesion volume distribution and correlation between lesion volume and the behavioral score**

**A:** Correlation between lesion volume and Center of Cancellation score. **B:** Lesion volume distribution over all 203 patients.

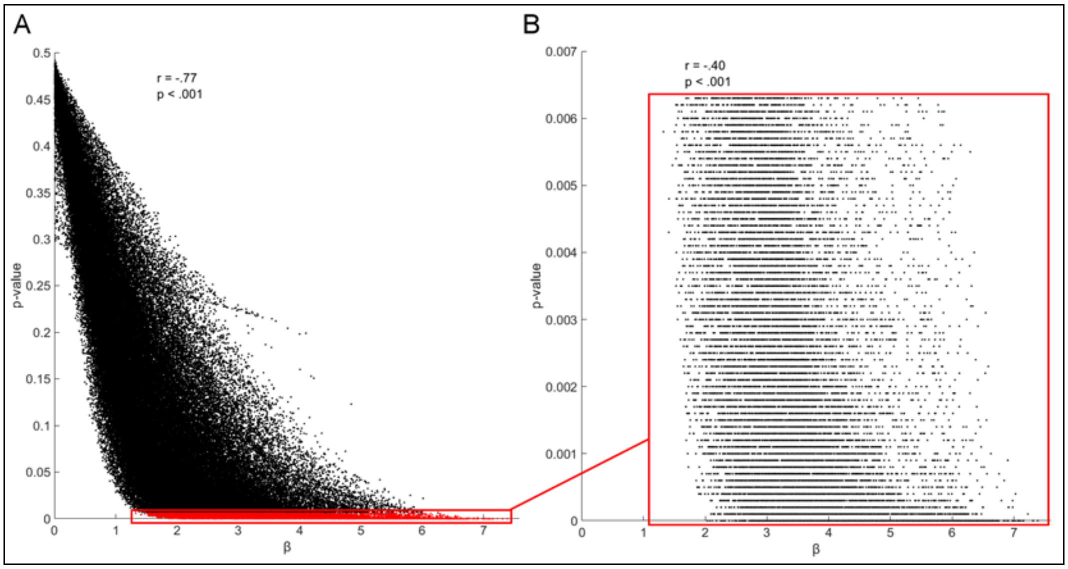


**Figure S4: Correlation between β-parameters and p-values of the main SVR-LSM analysis**

Scatter plots showing the relationship between β-parameters and p-values after applying lesion volume control with dTLVC for **A:** all voxels in the tested area and **B:** only significant voxels, FDR-corrected at $q = 0.05$, corresponding to a threshold of $p < 0.0063$. Demonstration that low β-parameters can yield low p-values and vice-versa.
